# Phylogenomic study of *Staphylococcus aureus* and *Staphylococcus haemolyticus* clinical isolates from Egypt

**DOI:** 10.1101/2021.05.01.442118

**Authors:** Cesar Montelongo, Carine R. Mores, Catherine Putonti, Alan J. Wolfe, Alaa Abouelfetouh

## Abstract

Antibiotic resistant *Staphylococcus* infections are a global concern, with increasing cases of resistant *Staphylococcus aureus* and *Staphylococcus haemolyticus* found circulating in the Middle East. While extensive surveys have described the prevalence of resistant infections in Europe, Asia, and North America, the population structure of resistant staphylococcal Middle Eastern clinical isolates is poorly characterized. We performed whole genome sequencing of 56 *S. aureus* and 10 *S. haemolyticus* isolates from Alexandria Main University Hospital. Supplemented with additional publicly available genomes from the region (34 *S. aureus* and 6 *S. haemolyticus*), we present the largest genomic study of staphylococcal Middle Eastern isolates. These genomes include 20 *S. aureus* multilocus sequence typing (MLST) types and 9 *S. haemolyticus* MLSTs, including 3 and 1 new MLSTs, respectively. Phylogenomic analyses of each species core genome largely mirrored MLSTs, irrespective of geographical origin. The hospital-acquired *spa* t037/SCC*mec* III/MLST CC8 clone represented the largest clade, comprising 22% of *S. aureus* isolates. Similar to other regional genome surveys of *S. aureus*, the Middle Eastern isolates have an open pangenome, a strong indicator of gene exchange of virulence factors and antibiotic resistance genes with other reservoirs. We recommend stricter implementation of antibiotic stewardship and infection control plans in the region.

**Impact Statement:** Staphylococci are under-studied despite their prevalence within the Middle East. Methicillin-resistant *Staphylococcus aureus* (MRSA) is endemic to hospitals in this region, as are other antibiotic-resistant strains of *S. aureus* and *S. haemolyticus*. To provide insight into the strains currently in circulation within Egypt, we performed whole genome sequencing of 56 *S. aureus* and 10 *S. haemolyticus* isolates from Alexandria Main University Hospital (AMUH). Through analysis of these genomes, as well as other genomes of isolates from the Middle East, we were able to produce a more complete picture of the current diversity than traditional molecular typing strategies. Furthermore, the *S. haemolyticus* genome analyses provide the first insight into strains found in Egypt. Our analysis of resistance and virulence mechanisms carried by these strains provides invaluable insight into future plans of antibiotic stewardship and infection control within the region.

**Data Summary:** Raw sequencing reads and assembled genomes can be found at BioProject Accession number PRJNA648411 (https://www.ncbi.nlm.nih.gov/bioproject/PRJNA648411).

## Introduction

Staphylococci are a heterogenous group of commensal bacteria in humans with the potential to cause infections ^1^. Two staphylococcal species especially relevant to the clinical setting are *Staphylococcus aureus* and *Staphylococcus haemolyticus*. *S. aureus* is arguably the most clinically important staphylococcal species; the infections it can cause range from mild erythema to serious life-threatening ailments, including septicemia, pneumonia, and endocarditis ^2^. A difficulty in treating and controlling *S. aureus* stems from its prevalence and increasing resistance to clinically used antibiotics, resulting in it being one of the leading agents for nosocomial and community-acquired infections ^3,4^. *S. haemolyticus* is the second most common staphylococcal species isolated in human blood cultures and a prominent reservoir for antibiotic resistance genes, which can be shared with other Staphylococci, including *S. aureus* ^5–7^.

Epidemiological surveillance and profiling are key to managing Staphylococci ^8,9^. Historically, profiling of Staphylococci has relied on complementary molecular typing strategies, such as Multi Locus Sequence Typing (MLST), typing of hypervariable short repeats in Protein A (*spa*), subtyping elements in the cassette chromosome mec (SCC*mec*), and presence of the Panton-Valentine leucocidin (PVL) ^10^. MLST consists of comparing the sequence of specific housekeeping genes in bacteria; the strategy is effective at tracking a broad range of clones over a global area, but prior to whole-genome sequencing (WGS) this method was slow and expensive ^10^. *spa* typing complements MLST by tracking the molecular evolution of *S. aureus*, given the relevance of Protein A to the infectious process ^9^. SCC*mec* permits profiling of clinically relevant antibiotic resistances, including *mecA*, which results in methicillin-resistant *S. aureus* (MRSA) ^11,12^. PVL is an important cytotoxin in MRSA that is common in community-acquired MRSA (CA-MRSA), but uncommon in hospital-associated MRSA (HA-MRSA) ^11^. Traditionally, these profiling strategies have been a powerful means to type, trace, and manage Staphylococcal infections, but technical limitations curtail the usefulness of molecular typing in real time ^9,10^.

The increasing utility, speed, and inexpensiveness of WGS in the clinical setting is poised to immensely benefit Staphylococci profiling ^10,13–15^. WGS allows access to the entire staphylococcal genome, including sequence data for typing MLST, *spa*, SCC*mec*, and PVL. In addition, WGS allows us to study the phylogenomic lineage, core, and accessory genome of isolates from an infectious outbreak or a geographical area. A key question is how WGS analysis compares to traditional typing techniques. For example, there is evidence that phylogenomic data does not always agree with standard typing methods; skepticism also exists that WGS can reliably detect single nucleotide polymorphisms (SNPs) in sensitive genetic content ^10^. In contrast, studies have demonstrated that WGS can be used to type, discriminate, and cluster staphylococcal isolates for the purpose of outbreak control ^13–15^. WGS could be used to close the gap in staphylococcal management in regions that have not been extensively monitored, such as the Middle East and specifically Egypt.

The epidemiology of Staphylococci in non-European countries of the Mediterranean region is under-studied ^16^. Antibiotic resistance in *S. haemolyticus* has been identified in Middle Eastern countries, such as Turkey ^6^ and Egypt ^7^. There is evidence that MRSA is prevalent and endemic to hospitals in this region, with a median MRSA prevalence of 38% in Algeria, Cyprus, Egypt, Jordan, Lebanon, Malta, Morocco, Tunisia and Turkey ^17^. Broadly speaking, PVL prevalence is reported as low in some of these countries, indicating a predominance of HA-MRSA ^16,18,19^. Research into the lineage of Staphylococci in this region is urgent, as it would give us both a present and future assessment of staphylococcal epidemiology.

Generally, molecular typing and phylogeny data are limited from this region. Multiple isolates in Palestine were typed as ST22 with a minority typed as ST80-MRSA-IV and PVL-positive ^20^. In Jordan, genotyping of *S. aureus* isolates revealed that the majority were ST80-MRSA-IV ^21^. In Lebanon, the primary lineage was PVL-positive ST80-MRSA-IV followed by PVL-positive ST30-MSSA ^17^. In Algeria, it was reported that ST80-MRSA-IV was present in most neonates tested over an 18-month period, with a minority of these PVL-positive ^22^. Finally, for Egypt, it has been reported that the prevalent MLSTs are ST30, ST80, and a novel type, ST1010; PVL prevalence has been estimated at 19% ^23^. Enany *et al*. reported that the Egyptian ST80 lineage was different from the globally prevalent ST80, primarily due to a unique *spa* type and antimicrobial resistance ^23^.

Egypt presents a unique case-study for staphylococcal distribution in Arab countries ^24^. Egypt’s cultural and geographical placement may facilitate local Staphylococcal exposure to international lineages, both from the Middle East and elsewhere. The accessibility of WGS presents an opportunity to profile Staphylococci in Egypt and the rest of the Arab region in terms of gene marker typing, core genome, and phylogenomics. Prior to this study, there were limited genomic data of *S. aureus* and *S. haemolyticus* in this region.

Here, we report the phylogenetic and phylogenomic associations of 56 *S. aureus* and 10 *S. haemolyticus* isolates from Egypt and their relationship to 34 *S. aureus* and 6 *S. haemolyticus* isolates from Egypt, Kuwait, Lebanon, Tunisia, Palestine, United Arab Emirates, Morocco, and Sudan. WGS afforded insight into the lineage and genetic content of these two staphylococcal species, including type information historically obtained using molecular methods. Both the MLST and SCC*mec* type mirrored the core genome, indicating that WGS is a fast and accessible option for Staphylococcal profiling. We identified multiple MLST and clonal complexes in circulation in the region, including 3 new genotypes. Genome analysis indicated that *S. aureus* in Egypt has an open pangenome that includes virulence genes in both the core and accessory genomes. Surveillance and profiling of Staphylococci are key to infection control, and we have shown that WGS can be a valuable asset, especially in regions where Staphylococci have not been well studied, such as the Middle East.

## Results

*S. aureus* and *S. haemolyticus* isolates were collected from patients presenting to the Medical Microbiology Laboratory at AMUH between September and December 2015. Draft genomes for 66 of the clinical isolates were of high quality and were included in our analysis. These genomes included 56 *S. aureus* and 10 *S. haemolyticus* isolates, assembled on average into 71 and 126 contigs, respectively. Additional publicly available *S. aureus* and *S. haemolyticus* strains were identified and included in subsequent analyses: 34 *S. aureus* (from Egypt n=17; Kuwait n=5; Lebanon n=4; Tunisia, Palestine and United Arab Emirates n=2 each; Morocco and Sudan n=1 each) and 6 *S. haemolyticus* (all from Egypt). **Supplementary Table S1** lists the available metadata and presents the genome assembly statistics for *S. aureus* and *S. haemolyticus*. The *S. aureus* genomes were, on average, larger than *S. haemolyticus* genomes: 2.8 Mbp and 2.5 Mbp, respectively. Genome size and GC content were on par with other publicly available genomes. Each draft genome was annotated using NCBI’s PGAP, identifying an average of 2,839 and 2,495 coding sequences (CDS) for *S. aureus* and *S. haemolyticus*, respectively. The strains varied in their number of rRNA operons and tRNAs.

### Strain genotyping

The genomes represent varied MLSTs. The 16 *S. haemolyticus* isolates examined here belonged to nine MLSTs, including a new genotype ST-74 (strain 51) assigned as a result of this study, and an isolate of unknown ST (strain 7A). ST-3 was the most common amongst the isolates examined (n=4) (**Supplementary Table S2**). A total of 20 *S. aureus* MLSTs were identified, including three novel types ST-5860 (strain 48), ST-5861 (strain 2705404), and ST-5862 (strain 2705410); all three of these strains came from prior studies and were isolated from Egypt, Kuwait and Lebanon, respectively (**Supplementary Table S2**). Twelve different MLSTs were identified among the Egyptian isolates; ST-239 was the most prevalent (n=24), followed by ST-1 (n=19) and then ST-80 (n=12). Two of the AMUH *S. aureus* isolates, strains AA32 and AA35, could not be typed due to incomplete sequences.

*S. aureus* isolates could be categorized into seven clonal complexes (CC) (**Supplementary Table S2**), the largest being CC8 (n=26), consisting mainly of Egyptian isolates and one Moroccan isolate (strain 12480433). 12 strains were identified as ST-80, which does not belong to a clonal complex. 20 different *spa* types were identified in addition to 7 isolates that could not be typed. The predominant *spa* type was t037 (n=33), with all but one belonging to CC8; 30 of these 33 isolates belonged to SCC*mec* III. The next most frequent *spa* type was t127 (n=19), all belonging to CC1. **Table 1** summarizes these results.

**Table 1:** MLST clonal complexes, *spa* types, and SCC*mec* types among the *S. aureus* isolates.

| MLST CC | Strain Name | Geographical Origin | *spa* type | SCC*mec* type |
| --- | --- | --- | --- | --- |
| CC1 | 3 (A), 3 (B), 23, 6 (B), 43, AA51, AA67, AA77 | Egypt | t127 | N/D |
|  | R181, R180, AA1, AA78 | UAE, Egypt | t127 | N/D |
|  | 6 (A), AA69 | Egypt | t127 | N/D |
|  | AA59, AA65, AA68 | Egypt | t127 | predicted as MSSA |
|  | AA103, AA87 | Egypt | t127 | predicted as MSSA |
| CC15 | 15, 16 | Egypt | t094 | predicted as MSSA |
|  | 17 | Egypt | unk | predicted as MSSA |
| CC22 | 41 | Egypt | t13828 | IV |
|  | Gaza_MRSA_B62 | Palestine | t223 | IV |
|  | AA18 | Egypt | t223 | IV |
|  | AA5 | Egypt | t3243 | IV |
|  | Gaza_MRSA_B04 | Palestine | t790 | IV |
|  | 40 | Egypt | unk | IV |
| CC30 | AA41 | Egypt | t037 | predicted as MSSA |
|  | 19 | Egypt | t1504 | N/D |
| **CC5** | AA30 | Egypt | t304 | IV |
|  | 14, AA76, AA80 | Egypt | t688 | N/D |
|  | AA70 | Egypt | t688 | VI(4B) |
| **CC8** | 12480433 | Morocco | t008 | IV |
|  | LHI_Sa_30 | Egypt | t008 | N/D |
|  | 46 | Egypt | t030 | III |
|  | 50, AA101, AA13, AA14, AA22, AA23, AA27, AA31, AA33, AA46, AA52, AA55, AA57, AA60, AA61, AA62, AA63, AA64, AA91, AA92 | Egypt | t037 | III |
|  | AA79 | Egypt | t037 | N/D |
|  | AA93 | Egypt | t037 | predicted as MSSA |
|  | AA29 | Egypt | unk | III |
| **CC97** | AA36 | Egypt | t267 | N/D |
|  | AA39, AA6 | Egypt | t267 | IV |
|  | AA104 | Egypt | t267 | N/D |
|  | AA8 | Egypt | unk | IV |
| **ST-80** | 2705432, 2705403, 2705405, 2705407, 2705409, 2705412 | Tunisia, | t044 | IV |
|  |  | Kuwait, |  |  |
|  |  | Lebanon |  |  |
|  | AA45 | Egypt | t044 | IV |
|  | 2705431 | Tunisia | t044 | predicted as MSSA |
|  | 2705411 | Lebanon | t131 | IV |
|  | AA2 | Egypt | t416 | IV |
|  | AA3, AA4 | Egypt | t416 | N/D |

### *Core and pangenomes of* S. haemolyticus *and* S. aureus *strains*

To investigate the core genome and pangenome of *S. haemolyticus*, we added the 6 publicly available *S. haemolyticus* genomes from Egypt to our 10 *S. haemolyticus* genomes (**Supplementary Table S1**). The pangenome for these strains included 3,541 genes (**Supplementary Fig. S1**), with 1,834 single copy number genes in the core genome. Included within these core genes are the virulence factors autolysin (*atl*), elastin binding protein (*ebp*), thermonuclease (*nuc*), and cytolysin (*cylR2*).

In addition to our 56 *S. aureus* genomes, our core and pangenome analysis included 34 *S. aureus* draft genome assemblies from the Arab region (**Supplementary Table S1**). The pangenome of these 90 isolates contained 4,283 genes (**Fig. 1, panel A**), the core genome included 1,501 single copy number genes, and the accessory genome contained 2,178 genes. These analyses show that the Arab isolates have an open pangenome. The functionality of the genes within the *S. aureus* core genome was determined according to their COG categories (**Fig. 1, panel B**). The core genome was further examined for virulence factors, finding the same gene related to autolysin (*atl*) that was found in the *S. haemolyticus* core. We also identified genes associated with intercellular adhesin, cysteine protease, thermonuclease, capsule, and the Type VII secretion system (**Table 2**).

**Table 2.** Virulence factors included in the *S. aureus* core genome.

| VFclass | Virulence factors | Related genes |
| --- | --- | --- |
| Adherence | Autolysin | *atl* |
|  | Intercellular adhesin | *icaA* |
|  |  | *icaD* |
|  |  | *icaR* |
| Enzyme | Cysteine protease | *sspC* |
|  | Thermonuclease | *nuc* |
| Immune evasion | Capsule | *cap5A* |
|  |  | *cap8B* |
|  |  | *cap5M* |
|  |  | *cap8N* |
|  |  | *capO* |
| Secretion system | Type VII secretion system | *esaB* |
|  |  | *essA* |
|  |  | *essB* |
|  |  | *esxA* |

**Figure 1.**
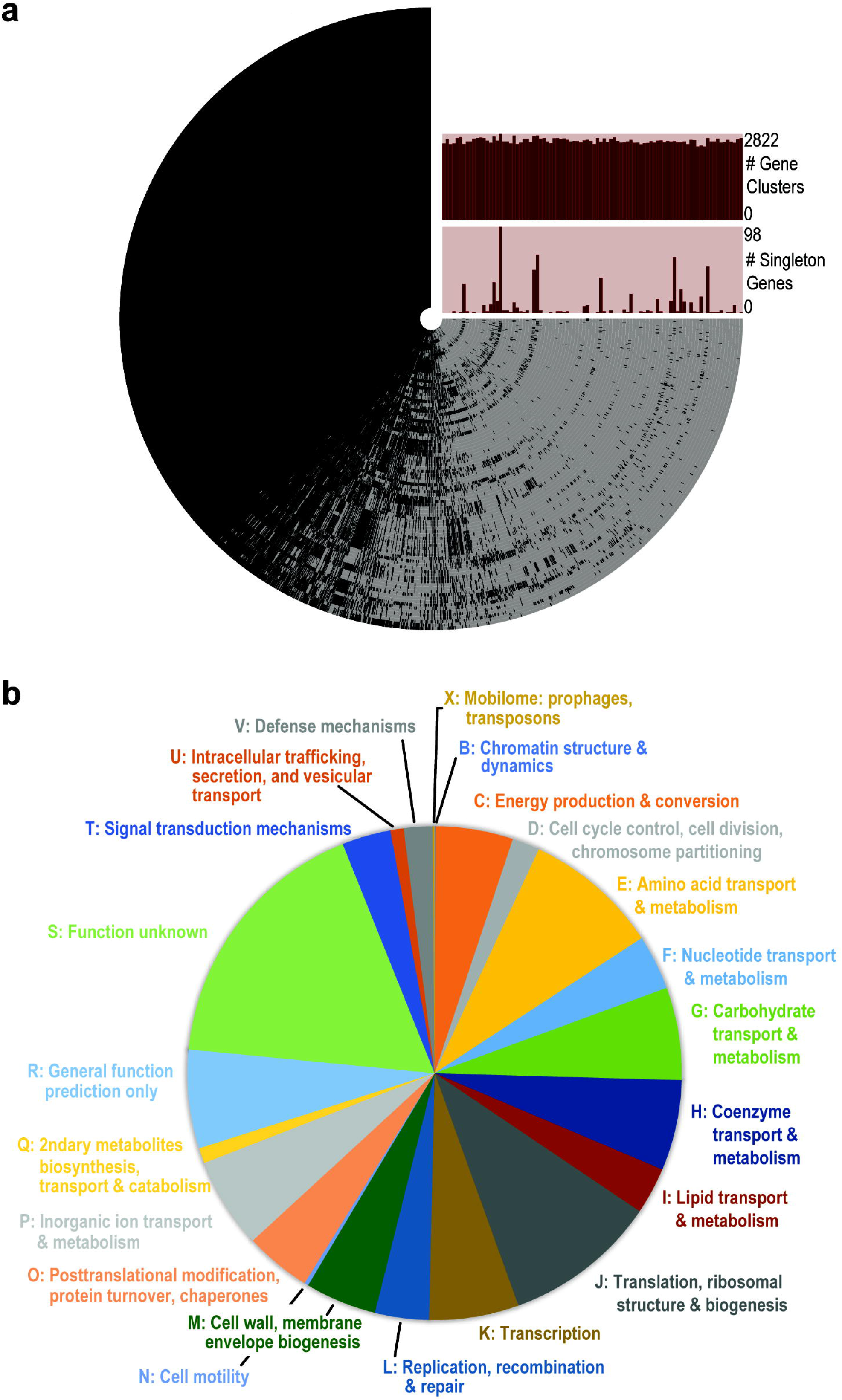
Genome analysis of 90 Arab *S. aureus* strains. **(a)** The pangenome. Each ring corresponds to a single genome. Each radial extension in the ring corresponds to the presence (black) or absence (light gray) of a given gene cluster (homologous gene). The bar charts list the number of genes identified in the given genome (top) and the number of singleton genes or genes that are unique to the given genome (bottom). The pangenome of these 90 isolates contained 4,283 genes, the core genome included 1,501 single copy number genes, and the accessory genome contained 2,178 genes. **(b)** Functionality of genes contained within the core genome. The same autolysin gene (*atl*) found in the core genome of *S. haemolyticus* was found in *S. aureus*.

In addition to the virulence factors found within the core genome, we identified virulence factors and antibiotic resistance genes within the accessory genome (**Supplementary Tables S3 and S4**). The isolates were screened for the presence of *lukF/S*-PV, which encodes PVL. Isolates positive for PVL were mainly (77%) *mecA* positive, present in CC1, ST-80, CC30 and CC8, and obtained from Egypt, Kuwait, Tunisia, Lebanon and Morocco. Importantly, isolates obtained from CA infections belonged to CC1, ST-80, CC5, CC97 and CC8, making PVL presence a good predictor for the ability of the isolate to cause CA infections.

### *Phylogenomic study of* S. haemolyticus *and* S. aureus *strains from the Arab region*

The core genes were used to derive phylogenies for each species. The *S. haemolyticus* isolates were all from Egypt and clustered into two clades (**Fig. 2**). As the tree shows, variation between the core genomes of these isolates was minor. Furthermore, the clade structure of the genomes corresponded with MLST, indicated in the bar of **Fig. 2**. The MLST tree for these genomes is shown in **Supplementary Fig. S2**.

**Figure 2.**
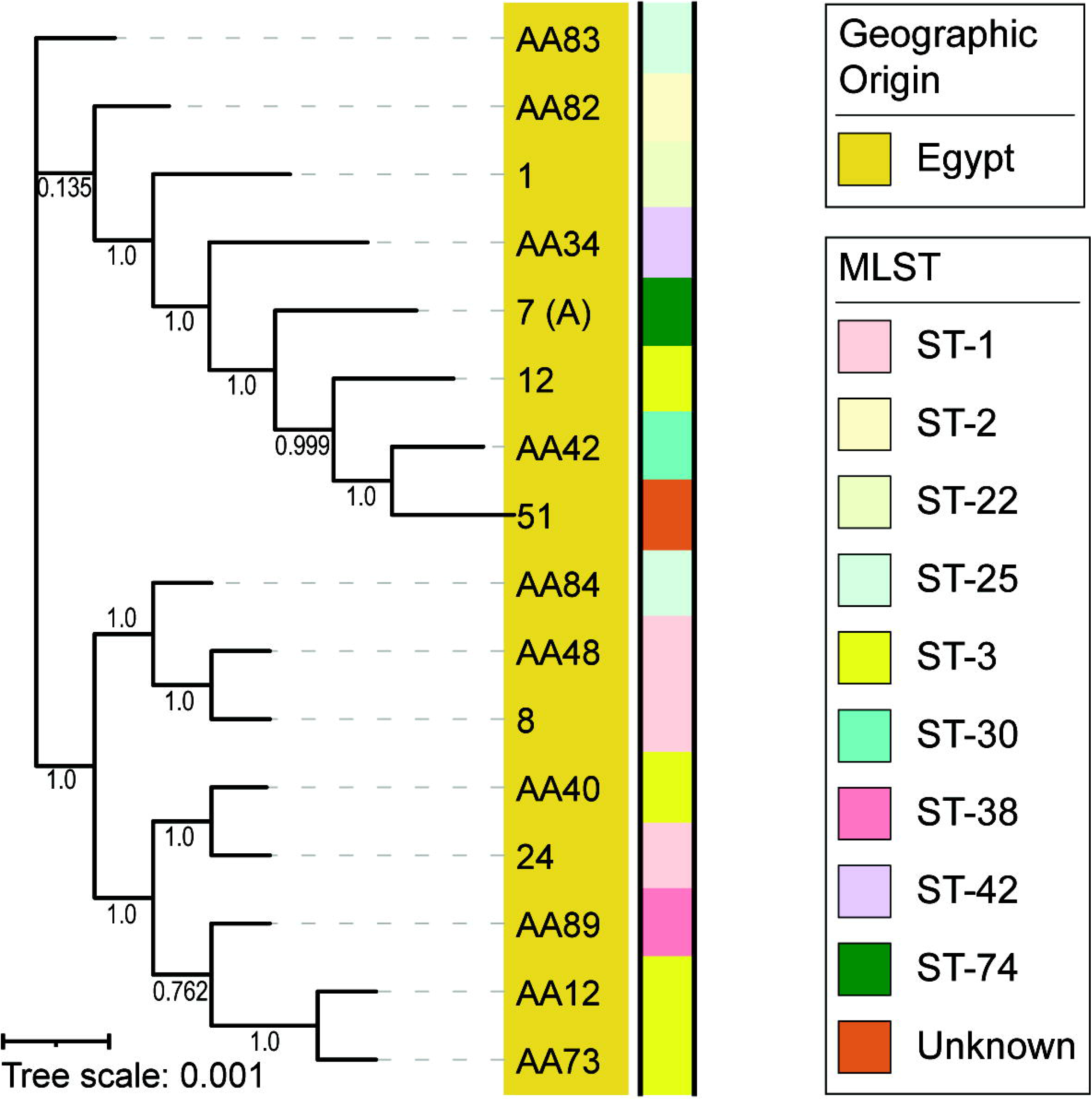
Phylogeny based upon the core genes for the *S. haemolyticus* isolates. All *S. haemolyticus* isolates were from Egypt and clustered into two clades corresponding with MLST.

*S. aureus* isolates came from all over the region, and clustered into six clades, with Egyptian isolates represented in all clades (**Fig. 3 and 4**). Clade 1 isolates belonged to ST-1 and were from Egypt and the UAE, clade 2 contained the majority of the Arab isolates including some from Egypt. The predominating clone seen among 46.7% of the isolates within clade 2 was *spa* t044/SCC*mec* IV/ST-80, which shows some degree of shared content between these isolates. Clade 3 isolates were solely from Egypt and belonged mainly to ST-15 and ST-5. Clade 4 comprised isolates from Egypt, Sudan and Palestine, with the majority belonging to ST-22 and ST-361. Clade 5 contained isolates from Egypt, belonging mainly to ST-97. The remaining isolates were in clade 6, of ST-239 and from Egypt, with the exception of one Moroccan isolate. This clade represents a *spa* t037/SCC*mec* III/MLST CC8 clone. The phylogenetic tree derived from the core genome sequences corresponded with the tree derived from the MLST marker genes (**Fig. 3** and **Supplementary Fig. S2**). 14 isolates lacked *mecA* (**Fig. 4**, pale green star) and occurred predominantly in CC1 (n=5), CC15 (n=3) and CC30, CC8 and ST-80, with one isolate in each; in addition, three isolates belonged to ST-361 (n=2) or ST-5860 (n=1). 13 of these *mecA* negative isolates were from Egypt.

**Figure 3.**
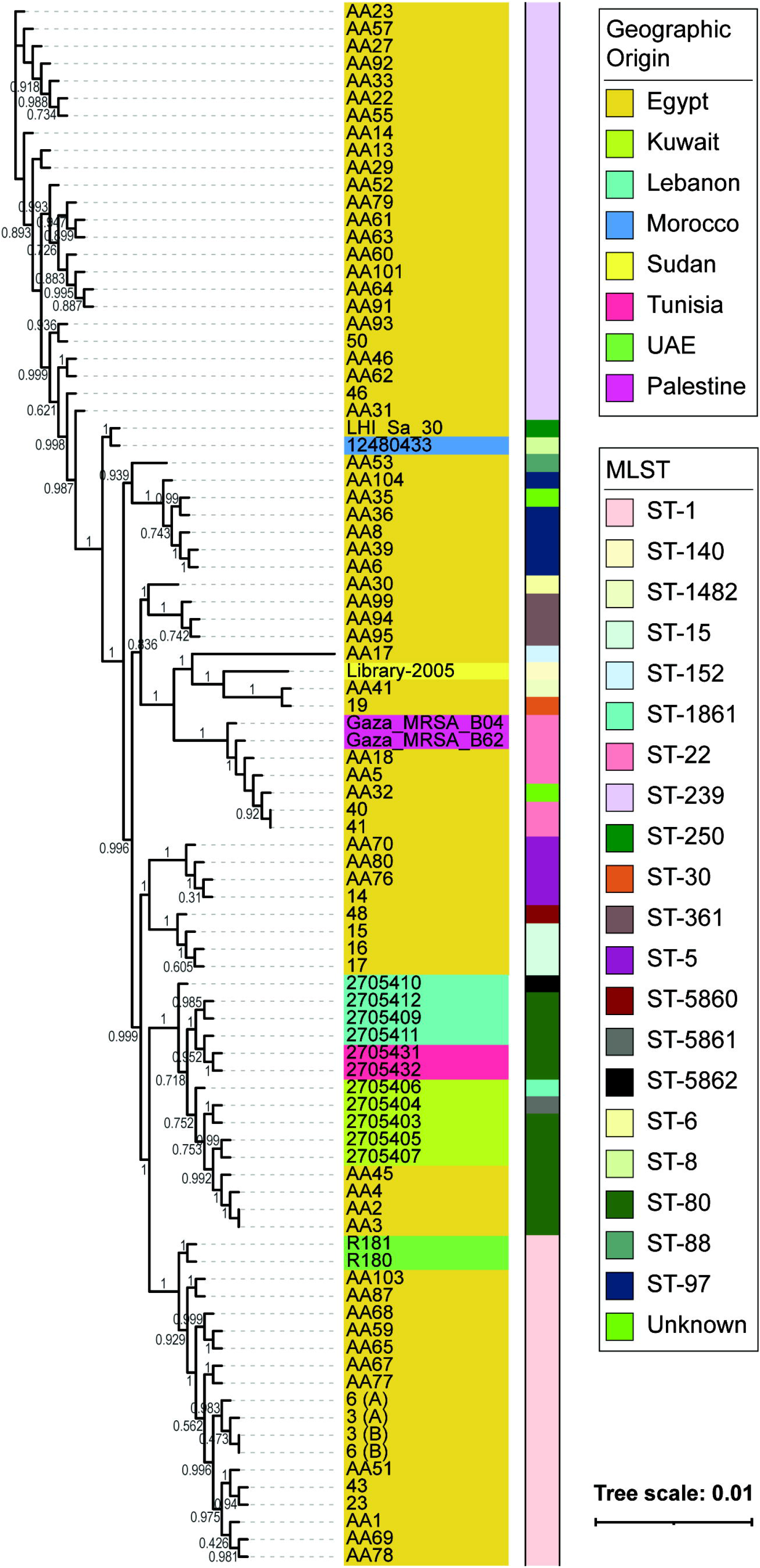
*S. aureus* core genome phylogeny colored by geographical origin of isolation (strain name color) and MLST (right bar). *S. aureus* isolates were from different parts of the region, and clustered into six clades, each containing Egyptian isolates. Clade 1 isolates belonged to ST-1 and were from Egypt and the UAE, clade 2 contained the majority of the Arab isolates, with *spa* t044/SCC*mec* IV/ST-80 as the predominating clone. Clade 3 isolates were solely from Egypt and belonged mainly to ST-15 and ST-5. Clade 4 comprised isolates from Egypt, Sudan and Palestine, with the majority belonging to ST-22 and ST-361. Clade 5 contained isolates from Egypt and belonged mainly to ST-97. The remaining isolates were in clade 6, of ST-239 and from Egypt and Morocco (n=1). This clade represents a *spa* t037/SCC*mec* III/MLST CC8 clone.

**Figure 4.**
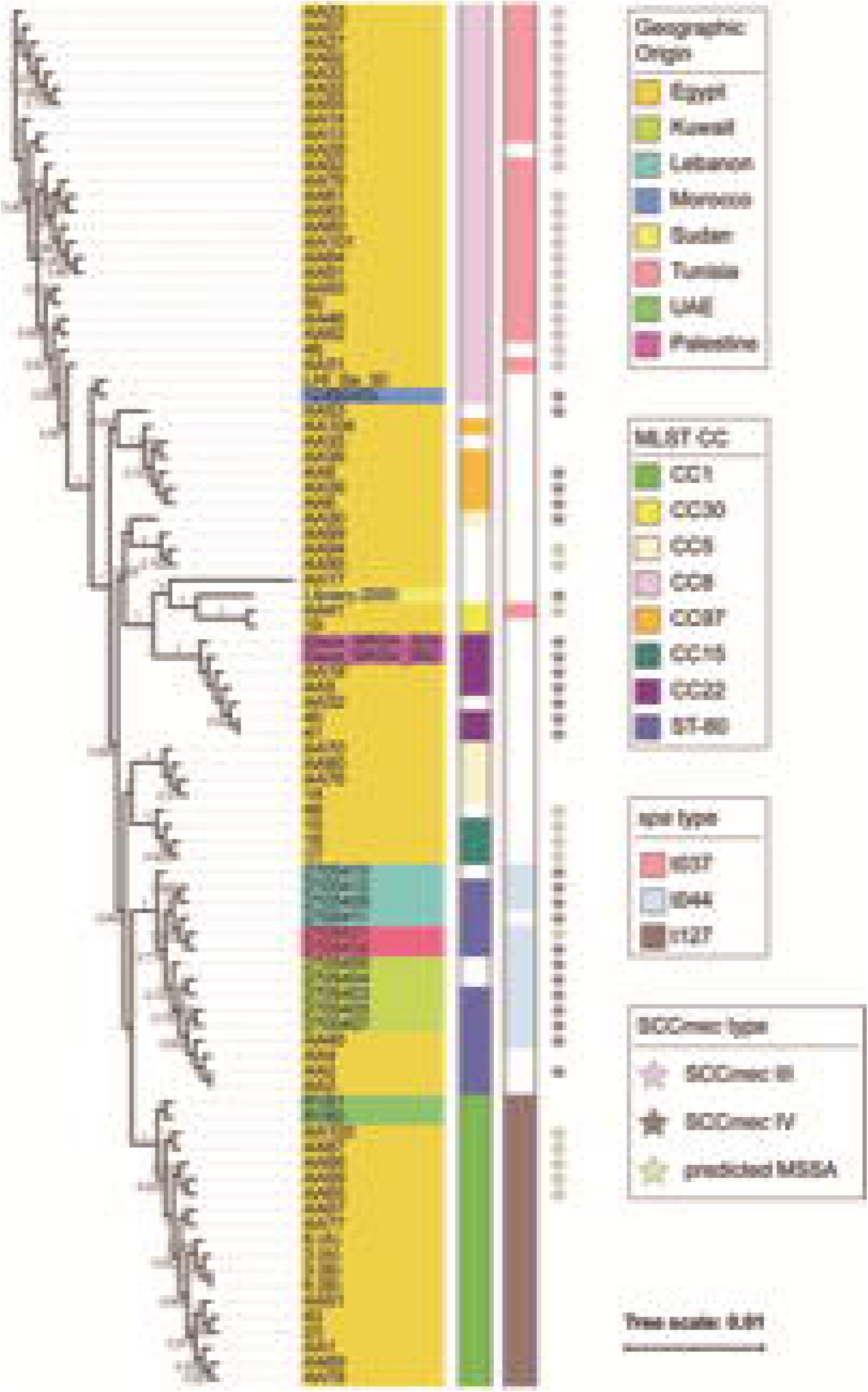
Phylogenetic tree based on core genome annotated by geographic origin, MLST CC, main *spa* and SCC*mec* types. 14 isolates, mostly from Egypt, lacked *mecA* and occurred predominantly in CC1 (n=5), CC15 (n=3) and CC30, CC8 and ST-80 (one isolate in each), ST-361 (n=2) or ST-5860 (n=1).

## Discussion

*S. aureus* is a major human pathogen in hospital and community settings, with the infection rate of MRSA increasing on a global scale at varying rates ^25,26^. While extensive surveys have provided insight into the prevalence of such resistant infections in Europe ^27^, Asia ^28,29^, and North America ^30,31^, limited data is available for the Arab region ^32^. Furthermore, antibiotic resistant *S. haemolyticus* strains have been identified worldwide ^5^, including Turkey ^6^ and Egypt ^7^. Prior to the study initiated here, there were limited genomic data for these two *Staphylococcus* species from the Middle East. The addition of 56 *S. aureus* and 10 *S. haemolyticus* genomes enabled our investigation of strain diversity within the region. With the majority (or all, in the case of *S. haemolyticus*) of genomes representing isolates from Egypt, we could investigate members of these species currently in circulation. We found that several different MLSTs and clonal complexes are in circulation within Egypt and more broadly within the region. 20 *S. aureus* MLSTs were identified in the region, including 3 new genotypes identified here, and 12 of these are in circulation within Egypt. Analysis of the *S. haemolyticus* genomes found 9 MLSTs in circulation within Egypt, including one new genotype.

The core genome for the *S. aureus* strains is slightly larger than that previously calculated for the species ^33,34^. This is expected, however, as our analysis is restricted to fewer genomes from a single region. Both our *S. haemolyticus* and *S. aureus* core genomes include the gene *atl*, which is essential for biofilm formation ^35^. Furthermore, the *ica* gene cluster, also associated with biofilm formation ^36^, as well as its regulator *icaR* ^37^, are included in the core genome of the *S. aureus* strains examined here. The presence of *atl* and the *ica* gene clusters signifies the biofilm potential of the isolates. This potential is relevant because 80% of human microbial infections are complicated by biofilm formation, such as in wounds, IV catheters, sutures and implants (see reviews ^38,39^). Moreover, the biofilm capacity to evade the host’s immune defenses, the inability of most antibiotic treatment regimens to eradicate existing biofilms and the fact that biofilms serve as a good medium for exchange of genetic material (e.g. plasmids) between cells make biofilm formation a major health concern in the clinical setting (^40^; see reviews ^41,42^).

The Arab *S. aureus* genomes have an open pangenome, evidence of gene exchange between these isolates and other reservoirs. *S. aureus* is naturally competent ^43^, and horizontal gene transfer (HGT) between strains, coagulase-negative *Staphylococcus* (CoNS) strains, and other species is well documented (see review ^44^). Recently, HGT was shown to be a driver of persistent *S. aureus* infections within patients ^45^. Genes within the accessory genome included virulence factors and antibiotic resistance genes (**Supplementary Tables S3 and S4**). Prior comparative genomic studies for this species similarly found an open pangenome and resistance genes within the accessory genome ^33^.

Phylogenomic analyses of the core genome largely mirrored MLST types (**Fig. 3**). This was irrespective of geographical origin. Interestingly, strains of the same SCC*mec* type had a more similar core genome sequence (**Fig. 4**). In a prior phylogenetic study, John and co-workers found that 16S rRNA gene sequence similarity did not correspond with SCC*mec* type, leading them to conclude that horizontal gene transfer plays a role in resistance gene acquisition ^33^. However, only two SCC*mec* types were identified for the samples examined here. Recently, Soliman *et al*. published a study characterizing the genomes of 18 MRSA isolates from a tertiary care hospital in Cairo, Egypt; their isolates were primarily SCC*mec* types V (n=9) and VI (n=2), not observed within our larger collection ^46^. Similarly, SCC*mec* type V and IV have been most frequently observed in other *S. aureus* studies within the region ^47–49^. Rather, our study found that SCC*mec* type III and IV were equally prevalent within the region. SCC*mec* type III predominated among HA-MRSA infections, suggesting that, in contrast to these prior studies, our isolates indicate that HA-infections have a higher incidence among the patients tested here. AMUH, where our isolates were collected, is the largest tertiary hospital and main referral center in the Northern sector of Egypt; thus, patients with more severe infections would be more likely to be treated at AMUH than at any other hospital in the region. Prior studies have found SCC*mec* type III to be the predominant type in Asian countries ^28^. Besides, the SCC*mec* type III/MLST ST-239 is the oldest pandemic strain of MRSA ^50^, which might explain its prevalence among the current collection of isolates.

The *S. haemolyticus* and *S. aureus* genomes examined here provide insight into the diversity of strains currently in circulation in Egypt, particularly with respect to their encoded virulence factors and antibiotic resistance genes. WGS analysis enabled a more complete picture of this diversity than molecular typing strategies. The *S. haemolyticus* genomes provide the first insight into strains found in Egypt. Identifying the main genotypes, as well as the resistance and virulence mechanisms among the resistant isolates in the region, can drive antibiotic stewardship and infection control plans.

## Methods

### Bacterial isolates

A total of 89 *S. aureus* and 14 *S. haemolyticus* consecutive non-duplicate isolates were collected from the Medical Microbiology Laboratory at Alexandria Main University Hospital (AMUH) between September and December 2015. These isolates were obtained from various clinical specimens, including pus, blood, sputum, urine, tissue, aspirate and broncho-alveolar lavage (BAL). The identity of the isolates was determined using conventional methods, such as Gram staining, growth on and fermentation of mannitol salt agar, growth on DNase agar and slide coagulase testing using Dryspot Staphytect Plus (Oxoid Ltd, England), and confirmed using Matrix Assisted Laser Desorption Ionization - Time of Flight Mass Spectrometry (MALDI-TOF MS) (Bruker Daltonik, USA). The isolates were further classified as either hospital-acquired or community-acquired infections based on a 48 h window between the dates of patient admission and isolate collection ^51^.

### DNA extraction

Colonies grown on tryptone soya agar (TSA) plates were harvested and washed in 1 ml phosphate buffer saline (PBS) and resuspended in 0.5 ml SET (75mM NaCl, 25mM EDTA, 20mM Tris, pH 7.5), to which 50ul of fresh 20 mg/ml lysozyme in PBS and 30ul Mutanolysin were added; the mixture was incubated at 37°C for 60 min. The cells were then treated with 60ul 10% sodium dodecyl sulphate and 20ul proteinase K and incubated at 55°C for two hours with gentle inversion. The suspension was mixed gently with 210ul of 6M NaCl, and 700ul phenol:chloroform were added, followed by incubation at room temperature for 30-60 minutes, using a rotating wheel for gentle mixing. The suspension was then centrifuged at maximum speed for 10 min and the aqueous phase was transferred to a new microfuge tube and mixed gently with an equal volume of isopropanol. The tubes were centrifuged to produce a DNA pellet that was washed with 70% ethanol, which was left to evaporate overnight. The pellets were resuspended in 50ul ddH_2_O and stored at −20°C till further processing.

### Genome Sequencing and Genome Assembly

The Illumina Nextera kit was used for whole genome library preparation. Each isolate was sequenced using the Illumina MiSeq System, producing paired-end 2×250 bp reads. Quality control and de-multiplexing of sequence data was done with onboard MiSeq Control software and MiSeq Reporter v3.1. Raw reads were trimmed using Sickle v1.33 (https://github.com/najoshi/sickle) and assembled using SPAdes v3.13.0 ^**52**^ with the “only-assembler” option for k◻=◻55, 77, 99, and 127. Genome coverage was calculated using BBMap v38.47 (https://sourceforge.net/projects/bbmap/). Contigs shorter than 500 bp were pruned using bioawk (https://github.com/lh3/bioawk). Genome assemblies were annotated using PATRIC v3.3.18 ^**53**^. Genomes were deposited in NCBI’s Assembly database, along with raw sequence data in SRA under BioProject PRJNA648411. Deposited genomes were annotated using the NCBI Prokaryotic Genome Annotation Pipeline (PGAP) v5.0 ^**54**^. Unless previously noted, default parameters were used for each software tool. To complement our analysis of the genomes from AMUH, raw sequence data for 41 *S. aureus* and 10 *S. haemolyticus* strains were retrieved from NCBI. These records were identified by searching SRA (as of January 2020) for strains isolated in the Arab region. These raw reads were processed as indicated above. High-quality assemblies were included in subsequent analyses.

### Bioinformatic Analysis

Multilocus sequence typing (MLST) was determined using the MLST v2.0.4 web server available through the Center for Genomic Epidemiology ^55^. MLST allele sequence and profile data were obtained from PubMLST v2.0.0 ^56^. *spa* typing was performed using the online tool SpaTyper v1.0 available through the Center for Genomic Epidemiology ^57^. SCC*mec* typing was performed using SCCmecFinder v1.2 online tool available through the Center for Genomic Epidemiology (https://cge.cbs.dtu.dk/services/SCCmecFinder/) ^58,59^. Resistance and virulence genes were identified using PATRIC v3.6.5 ^60^ and VFAnalyzer ^61^.

### Phylogenomic and Phylogenetic Analysis

The core and pangenomes were generated using anvi’o v5.1. The following scripts were used to calculate the pangenome: anvi-gen-genomes-storage, anvi-pan-genome and anvi-display-pan, and the following script was used to calculate the core genome: anvi-get-sequences-for-gene-clusters ^62,63^. Functional groups for the core genome were determined by querying core genome amino acid sequences against the COG database ^64^ through anvi’o. The core genes were concatenated for each genome and then aligned using MAFFT v7.388 ^65^. The tree was built using the FastTree v2 ^66^ plugin in Geneious Prime v2019.2 (Biomatters Ltd., Auckland, New Zealand). MLST ST sequences were downloaded from PubMLST v2.0.0 ^56^, aligned in Geneious Prime v2019.2 and the trees were built using the FastTree v2 ^66^ plugin in Geneious Prime v2019.2. iTOL v5.6.1 ^67^ was used to annotate and visualize all trees.

## Author Statements

## Acknowledgments

We acknowledge Roberto Limeira and Loyola Genomics Facility for performing the whole genome sequencing of the isolates. We also acknowledge funding from NIH (R01 DK104718 awarded to AJW), NSF (1661357 awarded to CP), USAID (GSP-T85 awarded to AA) and DFG (ZI 665/3-1 awarded to AA). The funders did not play a part in the design or conduct of the study

## Authors and Contributors

CM: Formal Analysis; Writing – Original Draft Preparation; Writing – Review and Editing CRM: Formal Analysis; Writing – Original Draft Preparation; Writing – Review and Editing CP: Formal Analysis; Writing – Original Draft Preparation; Visualization; Writing – Review and Editing

AJW: Formal Analysis; Conceptualisation; Writing – Review and Editing

AA: Conceptualisation; Formal Analysis; Writing – Original Draft Preparation; Writing – Review and Editing

All authors reviewed the manuscript.

## Data availability

Raw sequencing reads and assembled genomes can be found at BioProject Accession number PRJNA648411 (https://www.ncbi.nlm.nih.gov/bioproject/PRJNA648411)

## Ethics declarations

### Ethics approval and consent to participate

Not applicable

### Consent for publication

Not applicable

