## Supplemental Figures for "Phylogenomic study of *Staphylococcus aureus* and *Staphylococcus haemolyticus* clinical isolates from Egypt"

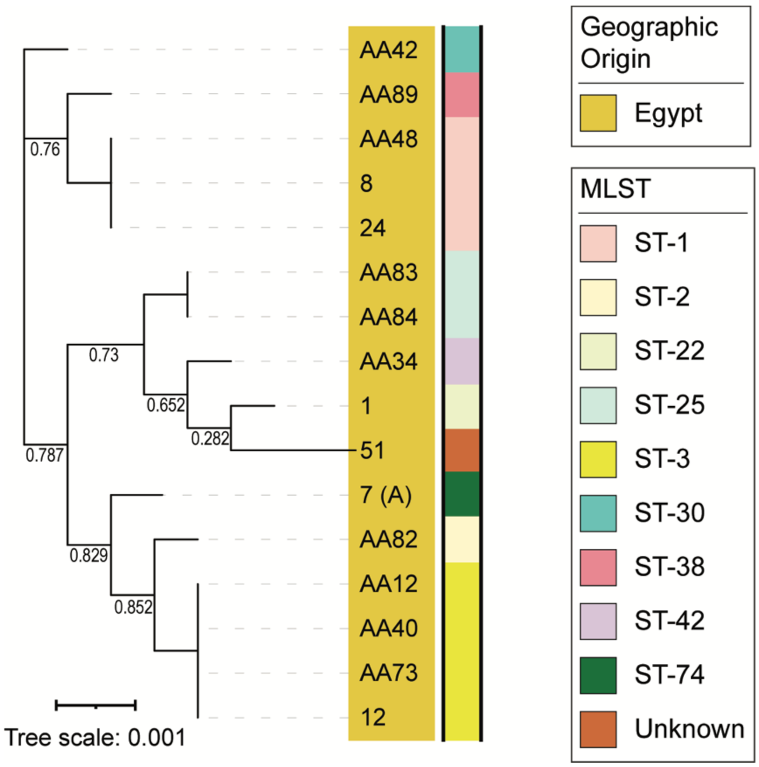


**Figure S1. MLST tree of *S. haemolyticus* annotated by geographic origin and MLST.**

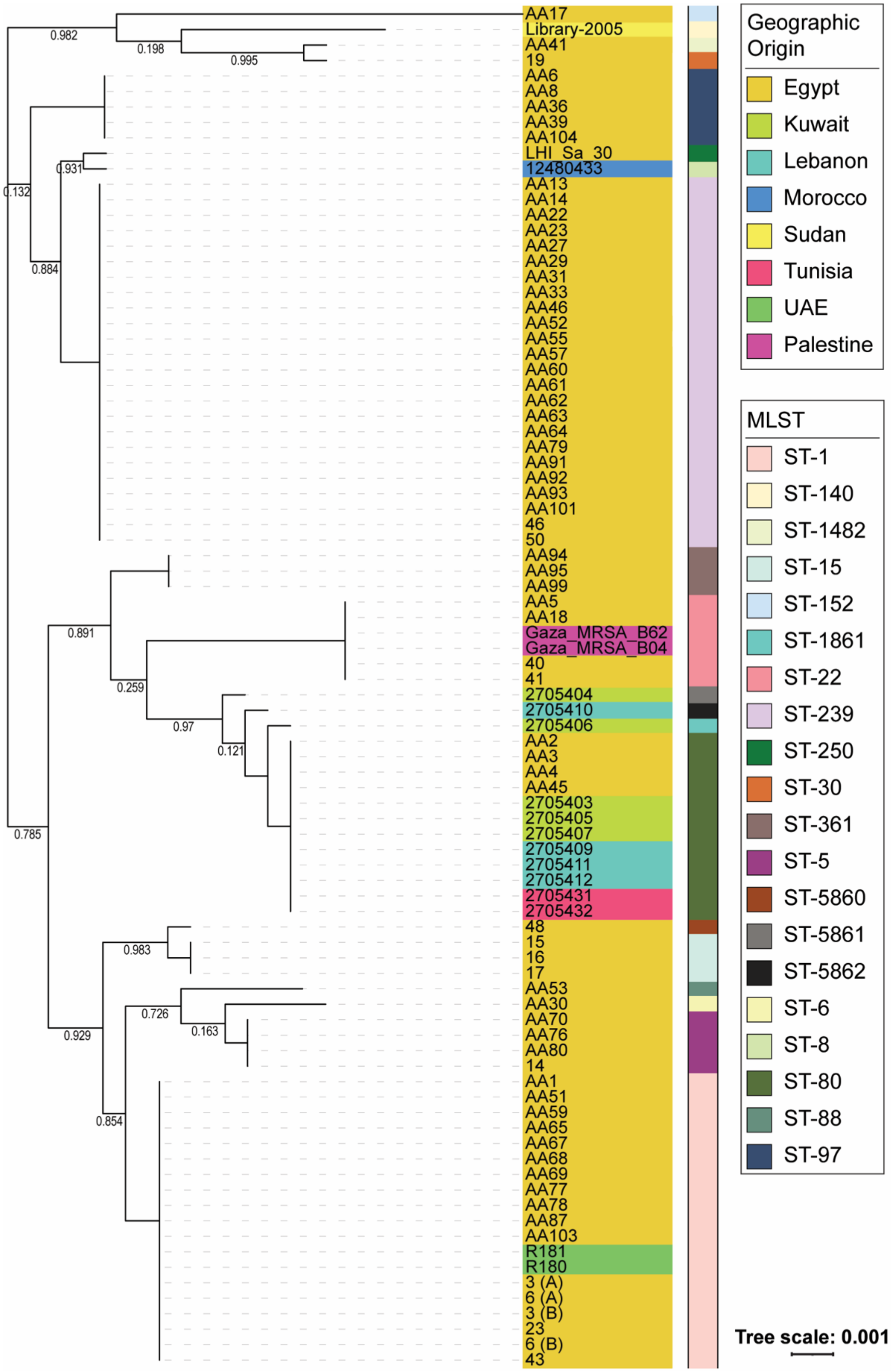


**Figure S2. MLST tree of *S. aureus* annotated by geographic origin and MLST.**
